# Divergent Pathogenic PR-DUB Complex Variants Converge Functionally Via PRC2 Displacement From Chromatin

**DOI:** 10.64898/2026.07.23.740410

**Authors:** Emma J. Doyle, Maeve Boyce, Sarah Buggle, Taylor Smith, Eugene Dillon, Anthony M. McElligott, Nina Orfali, Aisling Y. Coughlan, Diego Pasini, Eric Conway

## Abstract

The PR-DUB complex is responsible for erasing the repressive histone modification, H2AK119ub1. ASXL1-3 proteins are mutually exclusive catalytic partners of BAP1 in the PR-DUB complex. Somatic heterozygous *ASXL1-3* variants are associated with cancer, including myeloid malignancies, while *de novo* germline variants cause neurodevelopmental disorders such as Bohring-Opitz syndrome. These pathogenic variants are almost exclusively nonsense and frameshift and have been proposed to act as gain-of-function. However, the precise catalytic impact and mechanism of variant ASXL1-3 remains elusive. Using an isogenic embryonic stem cell model we have discovered that ASXL1 BOS variants drive reductions – but not global ablations – in H2AK119ub1, consistent with gain-of-function. This catalytic change occurs through the production of a truncated ASXL1 protein with enhanced stability. Hyper-stabilised ASXL1 drives a stoichiometric shift in PR-DUB assembly away from ASXL2 complexes. The drop in H2AK119ub1 levels ultimately reduces PRC2 binding and H3K27me3 deposition. Surprisingly, this phenotype is shared across PR-DUB loss-of-function models and indeed is emerging as a common phenotype across genetically and mechanistically distinct Polycomb-related chromatinopathies.

## Introduction

Polycomb Repressive De-ubiquitinase Complex (PR-DUB) is a multi-subunit chromatin regulatory complex that is responsible for the erasure of the repressive histone modification Histone H2A lysine 119 mono-ubiquitination (H2AK119ub1) (Sahtoe et al., 2016; Scheuermann et al., 2010). This catalytic function is performed through the core dimer of BAP1 and one of three obligate catalytic partners, ASXL1-3 (Scheuermann *et al*., 2010). Sequence variants in these catalytic dimer subunits are key drivers of human disease in both cancer and neurodevelopmental contexts (Bott et al., 2011; Doyle et al., 2022a; Tamburri et al., 2022; Testa et al., 2011). Monogenic heterozygous frameshift and nonsense variants in *ASXL1, ASXL2* and *ASXL3* are the defining features of the rare syndromes: Bohring-Opitz, Shashi-Pena and Bainbridge-Roper respectively (Bainbridge et al., 2013; Hoischen et al., 2011; Shashi et al., 2016). These variants are primarily *de novo* and have severe clinical presentations including intellectual disability, speech impairments, and seizures. In line with the critical role of PR-DUB as a chromatin regulator across distinct cell types during development (Dey et al., 2012; Fisher et al., 2010; McGrath et al., 2022; McGrath et al., 2021; Scheuermann *et al*., 2010), these symptoms affect multiple organ systems. This broad spectrum of clinical characteristics makes gene therapy design and delivery challenging. Such syndromes are part of a growing number of disorders defined by monogenic variants in chromatin regulators that have been termed ‘chromatinopathies’ or alternatively ‘mendelian disorders of epigenetic machinery’ (Fahrner and Bjornsson, 2019; Valencia and Pasca, 2022).

Similar, and overlapping, somatic variants in *ASXL1* have been identified in individuals with clonal haematopoiesis of indeterminate potential and in patients with overt myeloid haematological malignancies (Carbuccia et al., 2009; Gelsi-Boyer et al., 2009; Tamburri *et al*., 2022; Wang et al., 2021). Heterozygous frameshift ASXL1 variants alone can drive myelodysplastic syndromes in mice (Bai et al., 2021), albeit with incomplete penetrance. In patients, these variants represent an early clonal event in the development of myeloid malignancies and often co-occur with mutations in TET2, RUNX1, CEBPA and NRAS (Rose et al., 2017). Heterozygous frameshift variants in ASXL2 are also recurrent events in myeloid malignancies (Micol et al., 2014). These observations highlight how truncating variants in *ASXL1, ASXL2* and *ASXL3* have the capacity to cause human disease, with an affected individual’s phenotype depending on whether the variants are germline or somatic. Similarly, BAP1 is known to be a tumour suppressor due to loss-of-function variants having the capacity to drive solid tumours such as mesothelioma and uveal melanoma (Bott *et al*., 2011; Dey *et al*., 2012; Testa *et al*., 2011). Indeed, loss-of-function germline variants in key catalytic domains of BAP1 have been shown to define a distinct developmental disorder – Kury-Isidor syndrome (KIS) (Kury et al., 2022). Together, these *BAP1* and *ASXL1-3* variants underscore the importance of H2AK119ub1 equilibrium for typical development and maintenance of stem cell populations (Doyle et al., 2022b).

Mechanistically, the PR-DUB complex opposes the catalytic function of Polycomb Repressive Complex 1 (PRC1) - which deposits H2AK119ub1 through its catalytic subunits RING1A or RING1B (Scheuermann *et al*., 2010; Wang et al., 2004). H2AK119ub1 is primarily enriched at the promoters of developmentally repressed genes where it plays an instructive role in gene repression through downstream recruitment of Polycomb Repressive Complex 2 (PRC2) (Blackledge et al., 2014; Blackledge et al., 2020; Endoh et al., 2012; Tamburri et al., 2020). However, H2AK119ub1 also has emerging functions outside of these loci (Almeida et al., 2017; Fursova et al., 2019). Deposition of H2AK119ub1 in a broad ‘blanket’ distribution in intergenic regions is spatially constrained by the PR-DUB complex (Bonnet et al., 2022; Conway et al., 2021; Fursova et al., 2021). The catalytic activity of BAP1 maintains sufficient local enrichment of H2AK119ub1 at developmentally repressed promoters to recruit the PRC2 complex through its H2AK119ub1 readers - AEBP2 and JARID2 (Kalb et al., 2014). This facilitates discrete local distribution of the catalytic product of PRC2, H3K27me3, and the subsequent maintenance of plastic euchromatin (Conway *et al*., 2021). Therefore, despite PR-DUB opposing PRC1 catalytic function, it helps to maintain Polycomb repression through maintaining correct spatial distribution of H2AK119ub1 (Campagne et al., 2019; Kolovos et al., 2020) and H3K27me3 (Abdel-Wahab et al., 2012; Conway *et al*., 2021; Fursova *et al*., 2021). Structurally, ASXL1 binds to BAP1 through its N-terminal DEUBAD domain and is essential for PR-DUB catalytic activity (Sahtoe *et al*., 2016). It does so through directing the specificity of PR-DUB for its H2AK119ub1 substrate via direct contacts with DNA and the nucleosome acidic patch (Ge et al., 2023; Thomas et al., 2023). ASXL1 plays a further role in allosteric activation of DUB activity through ASXL1 mono-ubiquitination at lysine 351 within the DEUBAD. This acts as a molecular glue with the capacity to drive allosteric stimulation of H2AK119ub1 removal (Zhang et al., 2026).

While the BAP1 variants that are tumour suppressive in cancer and define Kury-Isidor syndrome are known to be loss-of-function (Conway *et al*., 2021; Fursova *et al*., 2021; Thomas *et al*., 2023), the impact of ASXL variants on H2AK119ub1 and the Polycomb system is not clear. Although functional studies have proposed that these truncating frameshift and nonsense variants may have a gain of deubiquitination activity (Balasubramani et al., 2015; Dong et al., 2025; Kohnke et al., 2024; Wang *et al*., 2021), these studies have not extensively examined the broader impact on Polycomb-mediated histone modifications, PR-DUB assembly, and gene repression. A number of mechanisms have been proposed for how ASXL variants may impact PR-DUB function, including stabilisation of BAP1, loss of FOXK1/2 interaction and gain of catalytic function (Kohnke *et al*., 2024; Wang *et al*., 2021; Xia et al., 2021). These are contrasted by studies suggesting that frameshift variants in *ASXL3* may lead to nonsense-mediated decay of mutant transcripts rather than production of functional ASXL3 protein (McGrath *et al*., 2022). To date, no study has systematically investigated these mechanisms in an isogenic setting to determine the impact of ASXL variants on PR-DUB composition, catalytic activity, and downstream impact on H2AK119ub1 and H3K27me3 genome-wide.

Here, we developed a tagged isogenic mouse embryonic stem cell (mESC) system to understand how *ASXL1* variants impact PR-DUB function, with a parallel established *BAP1* loss-of-function model as a contrast. These models allowed us to systematically characterise the impact of Bohring-Opitz syndrome (BOS) heterozygous G643Wfs on Polycomb repression and the epigenome. Critically, we found that frameshift variants dramatically stabilise ASXL1 which drives major protein level increases, whereas full length protein is inherently unstable. This increase in ASXL1 BOS truncation protein levels drives a global reduction in H2AK119ub1. Functionally, we show that this reduction in H2AK119ub1 is driven by a change in PR-DUB complex composition as ASXL1 BOS outcompetes ASXL2 for PR-DUB complex assembly. This is in stark contrast to BAP1 KIS models that have a global gain in H2AK119ub1. Strikingly, both BOS and KIS syndrome contexts converge through their impact on PRC2, with BAP1 KIS variants and ASXL1 BOS variants causing H3K27me3 and SUZ12 loss from chromatin, accompanied by dysregulation of developmental gene repression. Finally, examination of primary ASXL1-mutant AML and distinct PRC1 and PRC2-driven chromatinopathies reflect a strikingly similar phenotype of PRC2 activity loss. Taken together, this suggests that there may be a shared chromatin state underlying these genetically distinct diseases.

## Results

### Endogenous truncation variants cause stabilisation of ASXL1 through inhibition of proteasome-mediated degradation

To understand the clinical genetics underlying the rare PR-DUB syndrome-associated variants – Bohring-Opitz (*ASXL1*), Shashi-Pena (*ASXL2*), Bainbridge-Roper (*ASXL3*) and Kury-Isidor (*BAP1*) (Bainbridge *et al*., 2013; Hoischen *et al*., 2011; Kury *et al*., 2022; Shashi *et al*., 2016) – we collated data from published studies and mapped the reported variants according to their type and position (Supplementary Table 1). This meta-analysis of previous data highlighted the predominance of syndromic frameshift and nonsense mutations in the *ASXL* genes (Figure 1A). These variants are particularly enriched in the final two exons, a feature which is predicted to increase their chances of escaping nonsense mediated decay pathways that control the selective degradation of mRNAs with premature stop codons (Kim et al., 2024; Nakamura et al., 2026). Amongst the ASXL syndrome variants, the sole missense variant occurs within the Plant Homeodomain (PHD) of ASXL2 C1410G, highlighting a potentially critical role for this domain in ASXL-related syndrome pathogenesis. Previous studies have shown that the PHD domain of ASXL1-3 are critical for binding to MBD5 andMBD6 subunits of the PR-DUB complex and can contribute to stabilisation of ASXL proteins (Tsuboyama et al., 2022).

**Figure 1.**
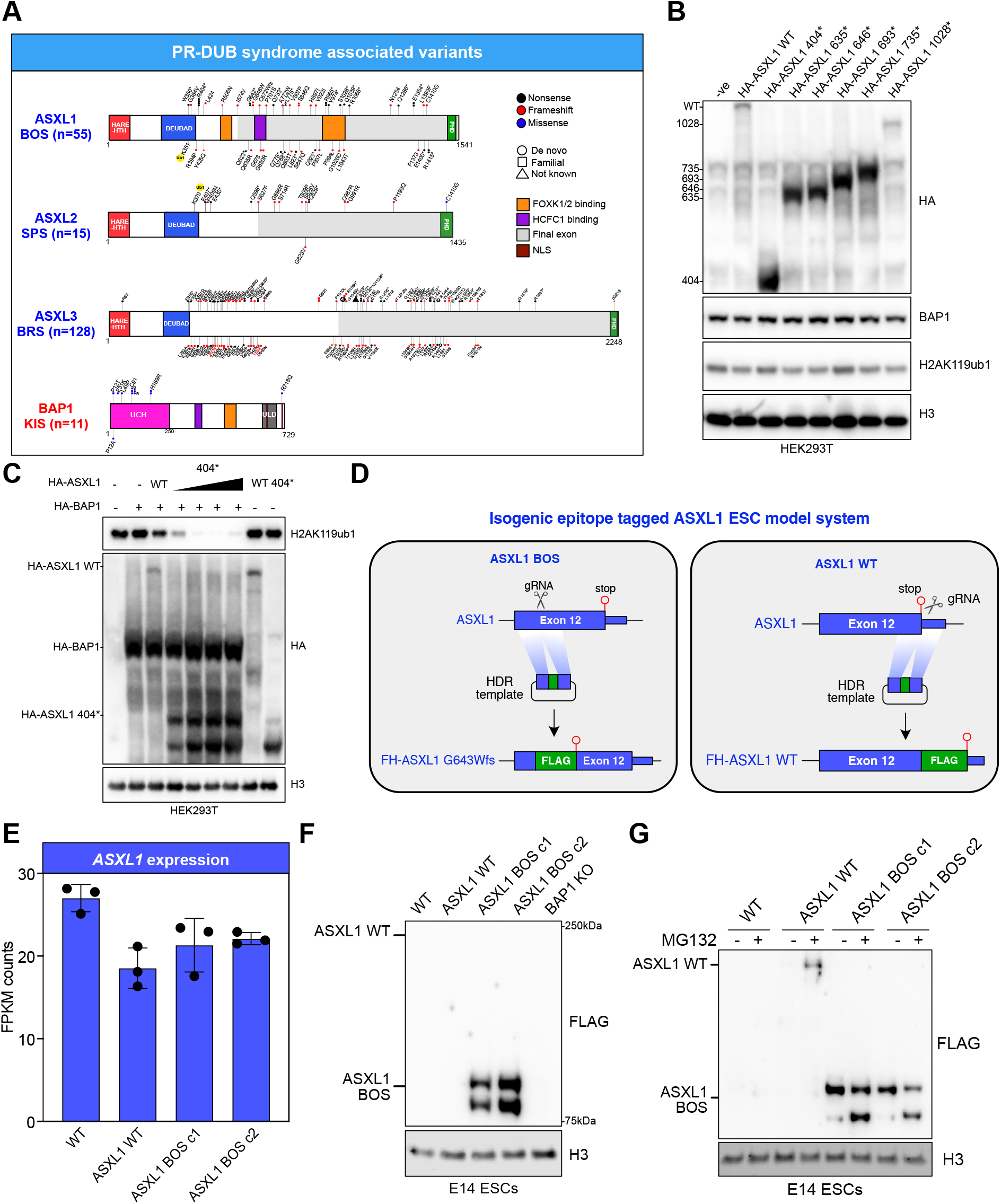
Endogenous truncation variants cause stabilisation of ASXL1 through inhibition of proteosome mediated degradation. **A**. Schematic of the human ASXL1-3 and BAP1 proteins summarizing the distribution of coding variants reported in Bohring-Opitz (ASXL1), Shashi-Pena (ASXL2), Bainbridge-Roper (ASXL3) and Kury-Isidor (BAP1) syndromes. Supplemental table S1 provides the source citations. Nonsense variants are shown in black, frameshift variants in red and missense variants in blue. Each shape represents a syndromic individual with a pathogenic PR-DUB variant that is either de novo (circle), familial (square) or of unknown inheritance status (triangle). **B**. Western blot with indicated antibodies on whole cell lysates of HEK293T cells following transient transfection with indicated plasmid construct. **C**. Western blot with indicated antibodies on whole cell lysates of HEK293T cells following transient transfection with indicated dose and pairing of plasmid constructs. **D**. Schematic of CRISPR nickase knock-in strategy to develop tagged heterozygous ASXL1FLAG-G643fs/+ (ASXL1 BOS) and ASXL1FLAG-WT/+ (ASXL1 WT) embryonic stem cells line. **E**. Quantification of ASXL1 transcript levels (FPKM) in indicated mESCs line from RNA-seq experiments. **F**. Western blot with indicated antibodies on whole cell lysates from the indicated mESCs line. **G**. Western blot with indicated antibodies on whole cell lysates from the indicated mESCs line treated with vehicle or MG132 for 6 hours.

On the other hand, Kury-Isidor syndrome variants in BAP1 were exclusively missense and are enriched in the catalytic UCH domain, with the exception being a missense variant in a nuclear localisation signal within the C-terminal extension (Kury *et al*., 2022). Based on their position in critical domains, these are considered loss-of-function variants, with missense variants at position C91 already having been established as catalytic null (Campagne *et al*., 2019; Conway *et al*., 2021; Scheuermann *et al*., 2010).

Due to the dual pathogenic functions of ASXL1 in Bohring-Opitz syndrome (BOS) and acute myeloid leukemia (AML) we chose to focus our study primarily on *ASXL1* variants. To understand the impact of *ASXL1* variants on PR-DUB and H2AK119ub1 we initially selected a panel of 6 nonsense variants and generated HA-tagged expression plasmids for each. Surprisingly, in HEK293T cells, exogenous expression of these variants alone had little impact on bulk H2AK119ub1 levels (Figure 1B), despite strong expression of the truncation variants relative to the full length wild-type ASXL1. Previous studies have established that exogenous co-expression of ASXL1 together with BAP1 is critical to drive global changes in H2AK119ub1 (Balasubramani *et al*., 2015). Therefore, we next co-expressed either ASXL1 wild-type (WT) or ASXL1 404* together with BAP1 in HEK293T. In keeping with findings from previous work, exogenous over-expression of BAP1 with ASXL1 404* led to dramatic global reductions in H2AK119ub1 levels, while the impact of co-expression of ASXL1 WT together with BAP1 was far milder (Figure 1C). However, we are unable to express the 1541 amino acid ASXL1 WT to the same extent as the truncated form – as detected by HA western blot – despite transfection of 7.5X more ASXL1 expression construct (lane 3 vs lane 4). While this could simply be due to complications with over-expression of a large protein, this led us to question whether the ASXL1 404* is more catalytically active or simply more abundant than ASXL1 WT. To address this, we established an isogenic mESC CRISPR knock-in system. We used CRISPR nickase to introduce a G643Wfs frameshift variant with in-frame FLAG/HA tags and subsequent stop codon (Figure 1D). Mouse *ASXL1* G643Wfs was selected as it corresponds to human ASXL1 G646Wfs which is the most frequently occurring ASXL1 variant in myeloid malignancies (Hoischen *et al*., 2011; Tamburri *et al*., 2022; Wang *et al*., 2021) and is found in 2, out of 55, known individuals with Bohring-Opitz syndrome (Figure 1A). This strategy, therefore, allows us to model the mechanism of ASXL1 dysfunction in both disease contexts. In parallel, we introduced FLAG and HA tags in frame with ASXL1 wild-type, immediately prior to the stop codon. As *ASXL1* variants tend to occur heterozygously in myeloid cancers and Bohring-Opitz syndrome, we selected heterozygous knock-in clones and extensively characterised and validated the genotype of both the knock-in allele (Figure S1A) and remaining wild-type allele. For simplicity, *ASXL1*^*FLAG-G643fs/+*^ and *ASXL1*^*FLAG-WT/+*^ embryonic stem cell models are hereafter referred to as ‘ASXL1 BOS’ and ‘ASXL1 WT’, respectively.

RNA-seq quantification of ASXL1 transcript levels (Figure 1E and Figure S1B) showed that the introduction of the G643Wfs variant did not lead to extensive nonsense mediated decay as there is only a minor, non-significant, change in *ASXL1* transcript levels in the knock-in clones. Our inclusion of epitope tags during CRISPR knock-in was in part driven by the absence of high quality antibodies generated against endogenous ASXL1 at the N-terminal portion that is shared in wild-type and truncated variants. FLAG western blot of the mESC models revealed substantial signal for the truncated form of ASXL1 in two distinct ASXL1 BOS clones (Figure 1F), however, it was difficult to detect the larger ∼177kDa full-length ASXL1.These low levels of ASXL1 WT reflect our original exogenous expression experiments in HEK293T (Figure 1B-C). Despite the low levels of ASXL1 WT, BAP1 endogenous co-immunoprecipitation showed that ASXL1 BOS and ASXL1 WT can still incorporate into the PR-DUB complex (Figure S1C).

This was expected due to the ASXL1 interaction region with BAP1 being located in the N-terminal DEUBAD domain (Sahtoe *et al*., 2016). To determine whether the observed low levels of ASXL1 WT are a consequence of reduced protein stability, we treated the mESCs with the proteasome inhibitor MG132 for 6 hours followed by western blotting. This experiment showed that ASXL1 WT is substantially stabilised by blocking proteasome function (Figure 1G), while ASXL1 BOS is only mildly stabilised. We observed a very similar phenotype in the exogenous HEK293T expression system wherein ASXL1 404*, 635*, 646*, 693*, and 733* nonsense variants exhibit a weaker fold change of protein levels after MG132 treatment compared to the full length ASXL1 and a nonsense variant that occurs further along the protein (ASXL1 1028*) (Figure S1D-F). This suggests that the truncated form of ASXL1 is indeed more stable. Recent reports suggest that there is a degron located within the central portion of ASXL1 that is lost in the truncated protein (Li et al., 2026), potentially explaining why truncated ASXL1 exhibits enhanced stability. Together these data show for the first time that endogenous nonsense and frameshift variant ASXL1 has dramatically increased stability through evasion of proteasome-mediated degradation.

### Physiological dosage of heterozygous ASXL1 frameshift variants cause genome-wide reductions in H2AK119ub1

To understand the impact that the stabilisation of truncated ASXL1 BOS has on PR-DUB catalytic activity in erasing H2AK119ub1 we examined the levels of this histone modification in the isogenic model system. Fluorescence western blotting (Figure 2A and B) revealed that H2AK119ub1 levels are indeed reduced in endogenous isogenic ASXL1 BOS cells, however, this is to a far weaker extent than in our, and previous, exogenous overexpression systems (Figure 1B) (Balasubramani *et al*., 2015). Quantification of H2AK119ub1 global levels showed that there is a 30-40% decrease in H2AK119ub1 levels across the two ASXL1 BOS clones. Importantly, endogenously tagging ASXL1 WT had no effect on H2AK119ub1 levels, while BAP1 knockout – which causes complete PR-DUB loss-of-function – resulted in a 2-3 fold increase in H2AK119ub1 in line with previous studies (Conway *et al*., 2021; Fursova *et al*., 2021). One proposed mechanism for gain in PR-DUB activity in the presence of truncated ASXL1 is increased stability of BAP1 itself (Wang *et al*., 2021). However, in this mESC system we see that BAP1 levels remain largely unchanged (Figure 2A and B). A recent study identified that mono-ubiquitination of K351 within the ASXL1 DEUBAD domain can drive allosteric activation of the complex and acts as a molecular glue for the catalytic PR-DUB dimer (Zhang *et al*., 2026). To understand whether K351 mono-ubiquitination contributes to the PR-DUB gain-of-function in the presence of truncated ASXL1, we co-expressed BAP1 with full length and 404* ASXL1 in HEK293T with and without K351R mutations (Figure S2A). These mutants maintain basic charge, but are an impaired substrate for mono-ubiquitination. This revealed clear loss of ASXL1 mono-ubiquitination, however lack of ASXL1 mono-ubiquitination did not impede the ASXL1 404* gain of catalytic activity, suggesting the change in this allosteric activity is not involved in gain of PR-DUB activity in ASXL1 BOS mESCs.

**Figure 2.**
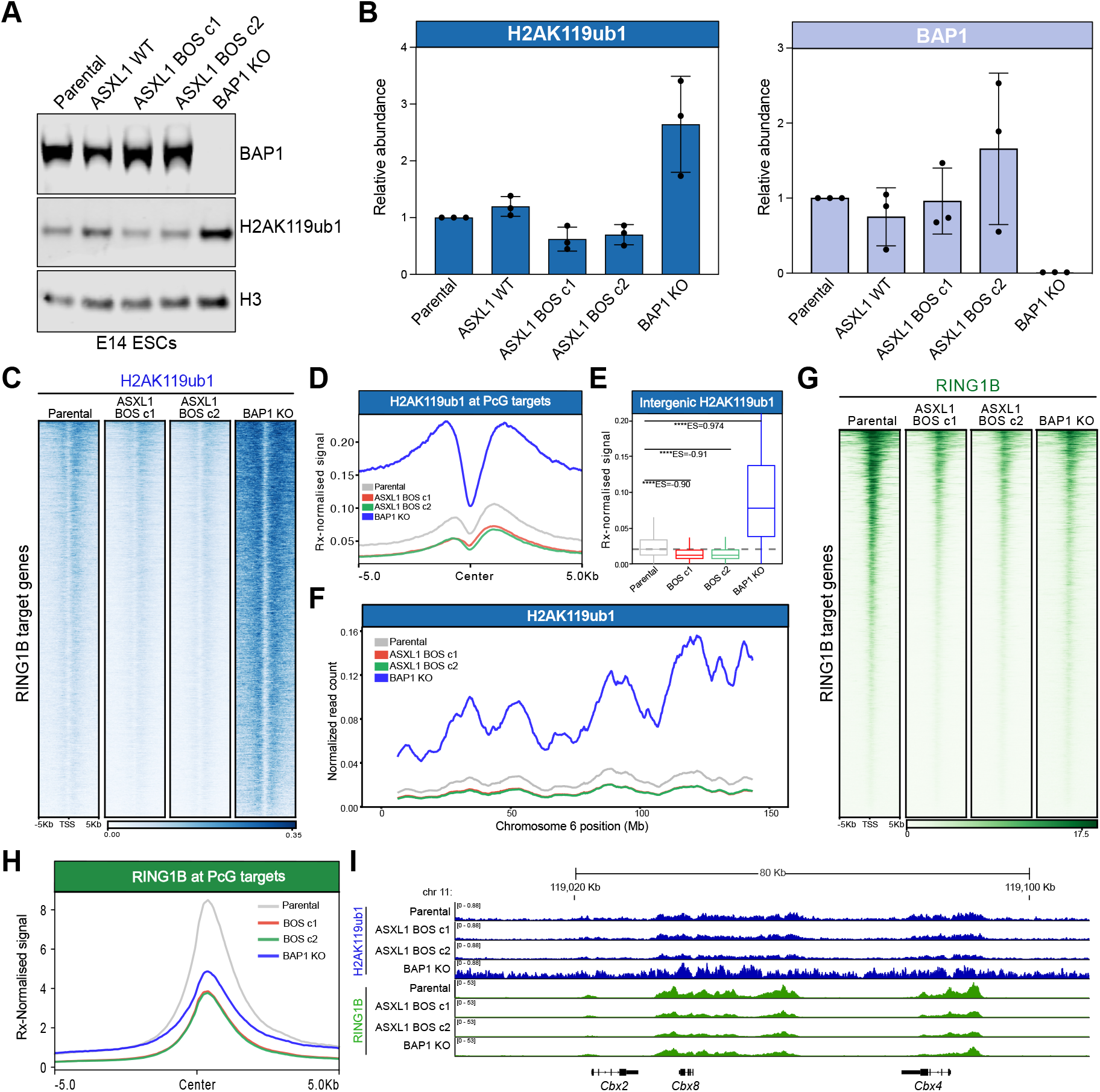
Physiological dosage of heterozygous ASXL1 frameshift variants cause genome-wide reductions in H2AK119ub1. **A**. Western blot with indicated antibodies on whole cell lysates from the indicated mESCs line. **B**. Quantification of fluorescence-based western blot in the indicated cell lines across three independent biological replicates. Signal intensity is normalised to H3 loading control and plotted relative to control cells. **C**. Heatmaps representing spike-in normalised ChIP-seq intensity for H2AK119ub1 at RING1B target genes (n=4,009) in the indicated cell lines. **D**. Average spike-in normalised ChIP-seq signal profile for H2AK119ub1 in the indicated cell lines at RING1B target genes. **E**. Boxplot representing H2AK119ub1 spike-in normalised ChIP-seq RPKM levels in the indicated cell lines at intergenic sites (n=38,239). **F**. Representation of the spike-in normalised read count of H2AK119ub1 in the indicated cell lines across chromosome 6 using 25kb windows. **G**. Heatmaps representing spike-in normalised ChIP-seq intensity for RING1B at RING1B target genes (n=4,009) in the indicated cell lines. **H**. Average spike-in normalised ChIP-seq signal profile for RING1B in the indicated cell lines at RING1B target genes. **I**. Genome browser snapshot of H2AK119ub1 and RING1B spike-in normalised ChIP-seq in indicated cell line.

As spatial distribution of histone modifications in the genome can be a more important determinant of their function than bulk levels (Brien et al., 2021; Conway *et al*., 2021), we next performed H2AK119ub1 spike-in normalised ChIP-seq. Examination of H2AK119ub1 levels at PRC1 (RING1B) target promoters (Figure 2C and D) showed that there is a mild loss of the repressive modification at these sites. Indeed, further analysis at intergenic regions and across entire chromosomes show this same modest, but consistent, reduction in H2AK119ub1 (Figure 2E and F). This is in line with the changes observed in bulk H2AK119ub1 levels (Figure 2A and B). These global reductions in H2AK119ub1 are in stark contrast to the major increases observed in BAP1 KO cells (Figure 2C-F). This is consistent with previous observations from ourselves and others that PR-DUB complex activity is not restricted to sites where it is stably bound (Bonnet *et al*., 2022; Conway *et al*., 2021; Fursova *et al*., 2021). Together, these parallel experiments show we have developed dual loss- and gain- of PR-DUB function isogenic model systems.

To define the impact of ASXL1 BOS variants on PRC1, we performed spike-in normalised ChIP-seq for RING1B. This revealed that there is a reduction in RING1B binding at PRC1 target genes in both the ASXL1 BOS and the BAP1 KO mESCs (Figure 2G-I). This observation is consistent with PRC1 catalytic mutant models that observed reductions in RING1B binding upon H2AK119ub1 loss (Blackledge *et al*., 2020; Tamburri *et al*., 2020). This could potentially be caused by disruption of the auto-regulatory feedback loop - driven by the capacity of RYBP to read H2AK119ub1 - that endows PRC1 with a dual read-write function (Ciapponi et al., 2024; Lopez et al., 2024; Zhao et al., 2020). This reduction in RING1B binding was a surprising common feature across the opposing gain- and loss-of-function models, suggesting some shared dysregulation of chromatin regulatory processes in distinct chromatinopathy contexts.

Overall, these data indicate that endogenous truncation of ASXL1 acts as a gain-of-function for PR-DUB catalytic activity – which is seemingly independent of BAP1 stabilisation or allosteric activation of ASXL1 – resulting in global reductions in H2AK119ub1 and downstream impacts on PRC1 chromatin occupancy.

### ASXL1 truncation drives a stoichiometric shift from ASXL2-PR-DUB to ASXL1-PR-DUB

While evident that the ASXL1 BOS containing complexes have increased deubiquitinase activity for H2AK119ub1, the molecular mechanism behind this catalytic change is unclear. To probe this question further, we examined the composition of PR-DUB. Studies to date have used exogenously tagged constructs to define the complex members (Campagne *et al*., 2019; Conway *et al*., 2021; Dong *et al*., 2025; Hauri et al., 2016; Kloet et al., 2016). However, due to the nature of these ectopic expression experiments, definition of complex subunit stoichiometry has remained evasive. Here we performed endogenous PR-DUB co-immunoprecipitation mass spectrometry for the first time (Figure 3A) in control mESCs to understand the relative abundances of each complex subunit. This stringent high salt protocol identified the majority of PR-DUB subunits expressed in mESCs (Figure 3B). The exception to this being FOXK1 and FOXK2 transcription factors, possibly suggesting their affinity for the complex is low. Crucially, defining the relative stoichiometries of the PR-DUB subunits in mESCs revealed that ASXL2 is the predominant paralog present in the complex in this context (Figure 3C), in line with previous reports from exogenous interactome studies (Kloet *et al*., 2016), while ASXL1 is only present in a minor fraction of PR-DUB complexes.

**Figure 3.**
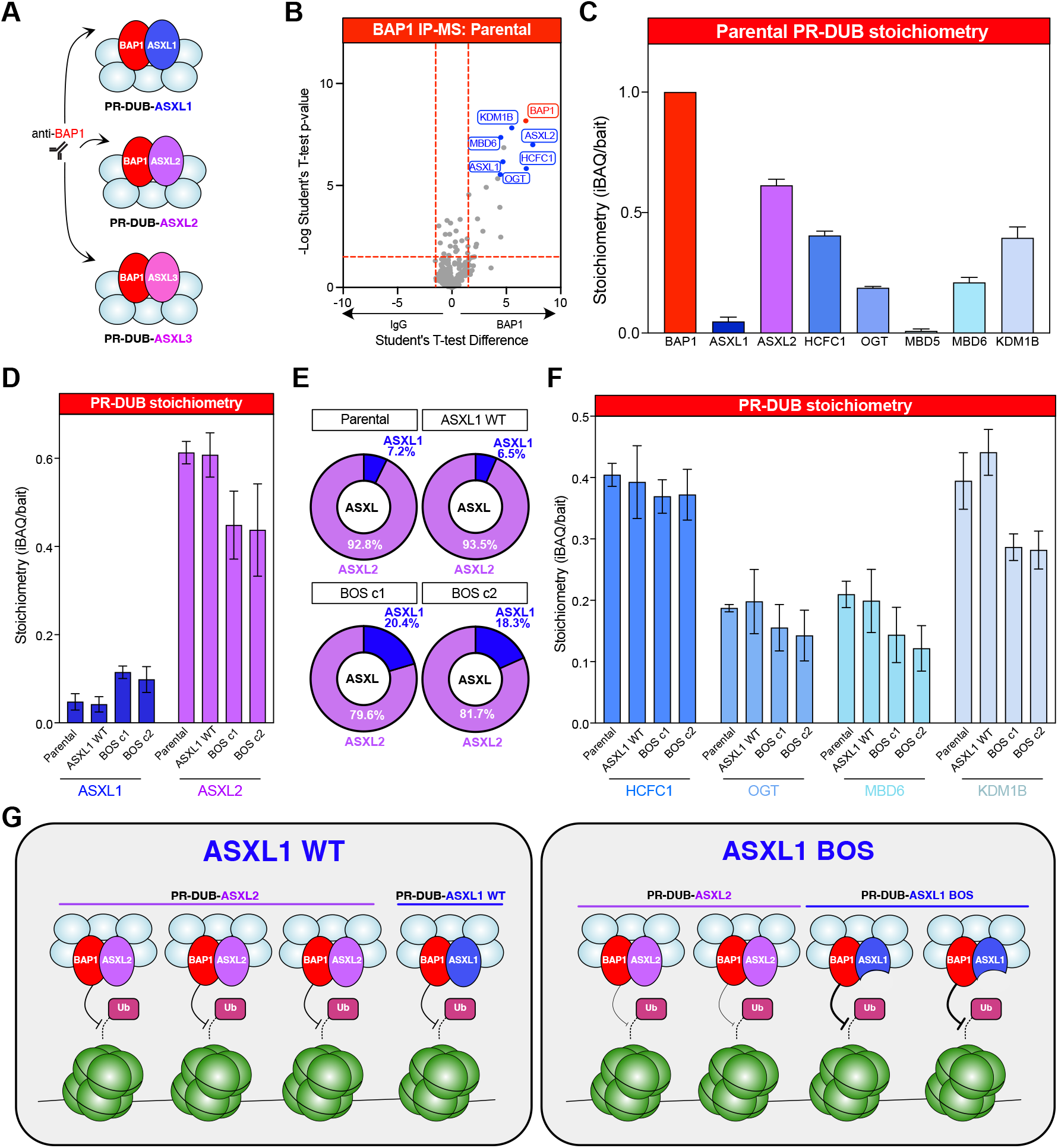
ASXL1 truncation drives a stoichiometric shift from ASXL2-PR-DUB to ASXL1-PR-DUB. **A**. Schematic of BAP1 endogenous co-IP mass spectrometry strategy to capture diverse PR-DUB complex assemblies. **B**. Volcano plot of endogenous BAP1 IP mass spectrometry data in wild-type mESCs relative to negative control IgG IP. T-test difference of LFQ values on the x-axis is plotted against -Log (T-test p-value). BAP1 (bait) is labelled in red and PR-DUB complex subunits are labelled in blue. **C**. Bar chart of stoichiometry (IBAQ relative to BAP1) of PR-DUB subunits in endogenous BAP1 co-IP experiments from wild-type mESCs. Data are represented as mean ±SD. **D**. Bar chart of stoichiometry (IBAQ relative to BAP1) of ASXL1 and ASXL2 in endogenous BAP1 co-IP experiments from the indicated cell lines. Data are represented as mean ±SD. **E**. Parts of a whole plots showing the relative percentage of ASXL proteins present in PR-DUB complexes in the indicated cell lines. **F**. Bar chart of stoichiometry (IBAQ relative to BAP1) of HCFC1, OGT, MBD6 and KDM1B in endogenous BAP1 co-IP experiments from the indicated cell lines. Data are represented as mean ±SD. **G**. Schematic showing the shift from ASXL2-PR-DUB complexes in WT ESCs to a blend of ASXL1- and ASXL2-PR-DUB complexes in ASXL1 BOS cell lines.

Expanding this PR-DUB composition analysis to the ASXL1 WT model revealed that endogenous tagging of ASXL1 WT did not affect complex composition or stoichiometry (Figure 3D-F and S3A-C). Importantly, the BAP1 interactome experiments in ASXL1 BOS mESCs showed a distinct shift in ASXL preference. Despite the heterozygous nature of the G643Wfs variant, the increase in stability of this protein was sufficient to drive an exchange of ASXL within the PR-DUB complex from 7.2% to 18.3-20.4% of complexes (Figure 3E). This increase in stoichiometry of ASXL1 is associated with a decrease of ASXL2 relative to BAP1 (Figure 3D) which suggests that there is competition between these paralogs for binding to BAP1. While little change is observed in HCFC1 and OGT assembly into PR-DUB, in the ASXL1 BOS mESCs there is a reduction in the levels of MBD6 that are present in BAP1 complexes (Figure 3F). This is in keeping with biochemical studies that have shown MBD6 and MBD5 bind to the C-terminal PHD domain of ASXL proteins (Tsuboyama *et al*., 2022), which is missing in the ASXL1 BOS truncation protein. In addition to this, there is a reduction in the levels of KDM1B identified within PR-DUB complexes in the ASXL1 BOS system, consistent with studies suggesting that KDM1B is exclusive to ASXL2-containing PR-DUB complexes (Campagne *et al*., 2019).

Taken together, these data suggest that the heterozygous BOS-associated truncation and stabilisation of ASXL1 causes a stoichiometric shift in PR-DUB complex assemblies that could contribute to its catalytic potency (Figure 3G). It is possible that variability in catalytic function between paralogs may contribute to this feature of PR-DUB complexes – as has been observed for other Polycomb systems – EZH2 having greater catalytic function than EZH1 for example (Lavarone et al., 2019; Margueron et al., 2008; McCole et al., 2025). It is tempting to speculate that this may also explain the relatively low number of Shashi-Pena (ASXL2) patients that have been diagnosed compared to the BOS (ASXL1) and Bainbridge-Roper syndrome (ASXL3) counterparts, which often feature more severe clinical presentations.

### BOS and KI variants converge via PRC2 eviction and H3K27me3 reductions

To investigate the downstream impact that PR-DUB gain-of-function has on PRC2, we next performed spike-in normalised ChIP-seq for SUZ12, a core component of PRC2 that is present in all sub-complexes, and H3K27me3, the repressive catalytic product of the complex. This revealed a remarkable impact of ASXL1 BOS variant PR-DUB on PRC2 chromatin occupancy SUZ12 binding is strongly disrupted at PRC1 target genes (Figure 4A-C). Interestingly, this impact is mirrored in the PR-DUB loss-of-function BAP1 KO model ESCs also, as we have shown previously (Conway *et al*., 2021), with SUZ12 binding at PRC1 targets being strongly decreased.

**Figure 4.**
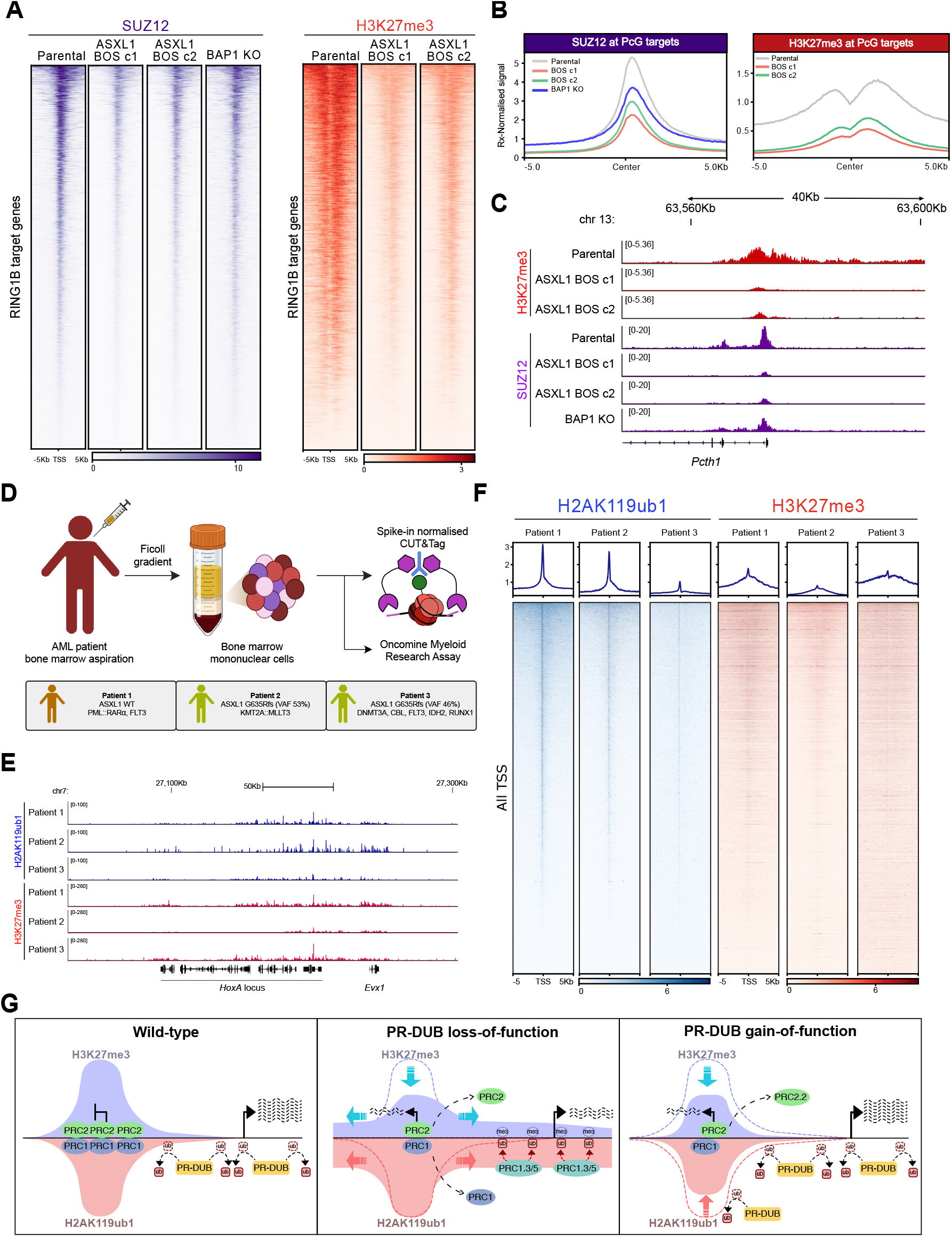
BOS and KI variants converge via loss of PRC2 and H3K27me3 reductions. **A**. Heatmaps representing spike-in normalised ChIP-seq intensity for SUZ12 and H3K27me3 at RING1B target genes (n=4,009) in the indicated cell lines. **B**. Average spike-in normalised ChIP-seq signal profile for SUZ12 and H3K27me3 in the indicated cell lines at RING1B target genes. **C**. Genome browser snapshot of H3K27me3 and SUZ12 spike-in normalised ChIP-seq in the indicated cell lines. **D**. Schematic of the biobanking and genotyping process for acute myeloid leukemia patient bone marrow mononuclear cells and subsequent spike-in CUT&Tag experiments. Patient mutations and cytogenetic information are summarised, with further detail in Table S2. **E**. Genome browser snapshot of H3K27me3 and H2AK119ub1 spike-in normalised CUT&Tag in the indicated patient BMNCs. **F**. Heatmaps representing spike-in normalised CUT&Tag intensity for H2AK119ub1 and H3K27me3 at all transcription start sites in the indicated patient BMNCs. **G**. Schematic of the proposed mechanism of action of ASXL1 truncating mutations in BOS and AML relative to the typical function of wild-type PR-DUB.

These changes in PRC2 occupancy extend to the catalytic deposition of H3K27me3 at Polycomb target sites in both PR-DUB loss- and gain-of-function models. Albeit, these concordant changes to PRC2 function seem to act through distinct mechanisms. As we and others have defined previously (Conway *et al*., 2021; Fursova *et al*., 2021; Zhang *et al*., 2026), PR-DUB loss causes global gains in H2AK119ub1, which in turn titrates PRC2 and its catalytic activity away from typical target loci to non-specific regions of the genome. However, the ASXL1 BOS gain-of-function model exhibits a divergent phenotype, as H3K27me3 activity appears to be reduced genome-wide (Figure 4A-C) in contrast to the redistribution of H3K27me3 observed in BAP1 KO ESCs (Conway *et al*., 2021).

While BOS is driven by variants that occur *de novo* in the germline (Hoischen *et al*., 2011), similar variants that occur in a somatic context can drive clonal haematopoiesis of indeterminate potential and myeloid cancers such as AML (Carbuccia *et al*., 2009; Gelsi-Boyer *et al*., 2009). We next assessed whether these somatic changes to *ASXL1* in an AML context re-shape the Polycomb repressive system in a similar way to ASXL1 BOS mESCs. Using primary bone marrow mononuclear cells (BMNC) from three patients with AML (Brophy et al., 2023), we performed spike-in normalised CUT&Tag for H2AK119ub1 and H3K27me3 (Figure 4D). The chromosomal and mutational profile of each patient performed as part of routine clinical diagnosis was known (Table S2) (Döhner et al., 2022). CUT&Tag of H2AK119ub1 in one ASXL1 wild-type patient (Patient 1) versus two ASXL1 G635Rfs mutant patients (Patient 2 – variant allele frequency 53% and Patient 3 – variant allele frequency 46%) revealed a consistent decrease in H2AK119ub1 at all promoters – in line with our observations in ASXL1 BOS mESCs (Figure 4E and F). In addition, we observed noteworthy reductions in H3K27me3 levels in one ASXL1 mutant sample (Patient 2) compared to ASXL1 wild-type (Figure 4E-F). No such reduction was observed in Patient 3, perhaps due in part to the presence of a co-occurring mutation in DNMT3A that may compensate for H3K27me3 deposition through indirect mechanisms relating to H3K36me2 (Streubel et al., 2018; Weinberg et al., 2021). Although correlative, this orthogonal approach in a distinct primary disease model showed that H2AK119ub1 and PRC2 activity can be disrupted in the context of PR-DUB gain-of-function.

This reduction of PRC2 binding and activity may occur through higher PR-DUB activity driving the lower levels of H2AK119ub1 (Figure 2C) at PRC1 targets, resulting in a weaker capacity to recruit PRC2.2 complexes, containing the H2AK119ub1 reader subunits AEBP2 and JARID2 (Figure 4G) (Blackledge *et al*., 2020; Kalb *et al*., 2014; Kasinath et al., 2021; Tamburri *et al*., 2020). The downstream consequences of this are reduced PRC2 at Polycomb targets, but without redistributed H2AK119ub1 to direct them elsewhere there is no intergenic accumulation of H3K27me3.

### Dysregulation in Polycomb target gene regulation underlies Polycomb-related syndromes

To further understand whether the disruption to PRC1 and PRC2 function in the ASXL1 BOS model translates to impacts in gene regulation, we performed RNA-seq. This revealed that the PR-DUB gain-of-function model has a distinct transcriptional phenotype compared to BAP1 KO with fewer significantly differentially expressed genes (Figure 5A and B). Both gain- and loss-of-function of PR-DUB models have a similar number of up- and down-regulated genes, suggesting that their impact is not as simple as complete loss of a repressor like RING1B (Tamburri *et al*., 2020), with potential indirect effects coming from the stable clonal nature of the ESC system. In fact, this milder transcriptional phenotype could explain why these germline heterozygous variants in *ASXL1* are observed in humans, while complete loss of PRC2 or PRC1 function are not observed. Based on mouse models, complete loss of PRC1 or PRC2 functions are early embryonic lethal (O’Carroll et al., 2001; Pasini et al., 2004; Voncken et al., 2003). Further analysing the commonly dysregulated genes across our BOS model mESCs revealed that developmental processes are highly enriched among the downregulated (anterior/posterior pattern specification, embryonic limb morphogenesis) and upregulated (kidney development) biological processes (Figure 5C and D, Figure S4D). The presence of these general developmental terms may be indicative of the incomplete disruption to the Polycomb system and a subsequent loss of the precise timing and scale of control of gene repression during cell fate decisions. Indeed, many of the dysregulated genes are PRC1 targets (Figure 5E). The disparate developmental processes impacted in the ASXL1 BOS cells certainly mirrors the pleiotropic developmental phenotypes displayed by people affected by ASXL-syndromes.

**Figure 5.**
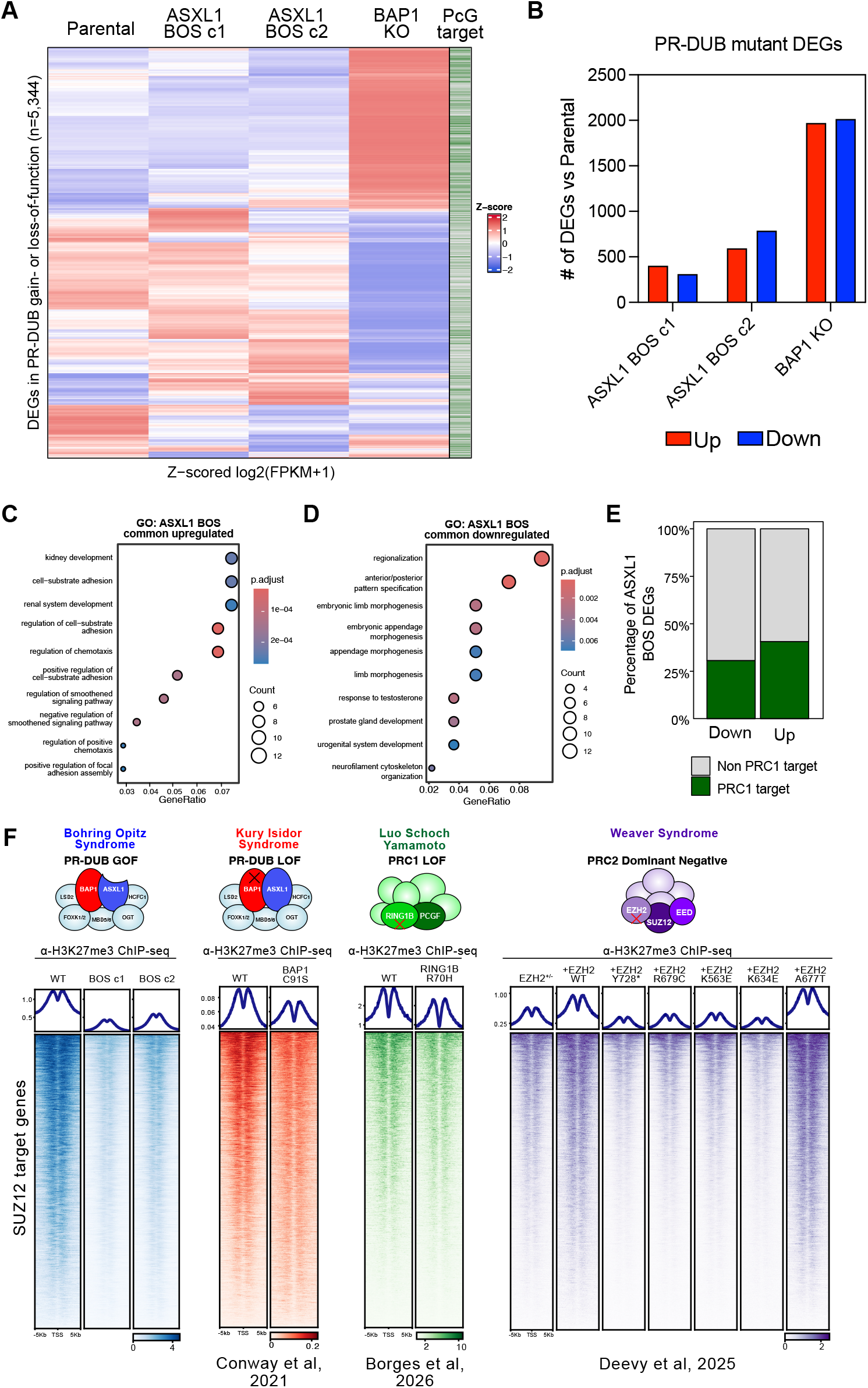
Dysregulation in Polycomb target gene regulation underlies Polycomb related syndromes. **A**. Heatmap of RNA-seq Z-scores for genes that are differentially expressed in any of ASXL1 BOS c1, c2 or BAP1 KO cell lines relative to wild-type following unsupervised hierarchical clustering. PRC1 target genes are labelled green in the right hand column. **B**. Bar chart illustrating the number of significantly differentially expressed genes in the indicated cell lines compared to WT mESCs. Genes were considered differentially expressed when presenting a log2 fold change equal to or greater than 1 and an adjusted p-value lower than 0.05. **C**. Gene ontology analysis of biological processes enriched in genes downregulated in both ASXL1 BOS clones. **D**. Gene ontology analysis of biological processes enriched in genes upregulated in both ASXL1 BOS clones. **E**. Parts of a whole plot showing the percentage of DEGs from both ASXL1 BOS clones that overlap with PRC1 complex target genes. **F**. Top. Schematic of PR-DUB, PRC1 and PRC2 complexes indicating the genetic aetiology of their associated syndromes. Bottom. Heatmaps representing ChIP-seq intensity for H3K27me3 at SUZ12 target genes (n=5,014) in the indicated cell lines from studies on PR-DUB, PRC2 and PRC1-related chromatinopathies (Borges *et al*., 2026; Conway *et al*., 2021; Deevy *et al*., 2025).

The number of identified chromatinopathies driven by monogenic events in chromatin regulator coding genes is growing as sequencing for patients with rare diseases becomes more accessible (Fahrner and Bjornsson, 2019; Valencia and Pasca, 2022). Aside from ASXL-syndromes, many chromatinopathies have been defined that impact Polycomb Repressive Complex subunit coding genes. There has been a flurry of recent mechanistic studies on these syndromes, including on Weaver (EZH2), Kury-Isidor (BAP1) and Luo-Schoch-Yamato (RING1B) (Borges et al., 2026; Conway *et al*., 2021; Deevy et al., 2025). Despite contrasting genetic mechanisms – including gain-of-function, dominant negativity and loss-of-function – each of these studies have observed major detrimental impacts of syndrome-associated variants in isogenic mESC model systems. To put our results in the context of the growing number of Polycomb-related chromatinopathies, we analysed the respective H3K27me3 ChIP-seq datasets at PRC2 target genes and found striking similarities in both the direction and extent of impact on H3K27me3 levels at these target loci (Figure 5F) (Borges *et al*., 2026; Conway *et al*., 2021; Deevy *et al*., 2025). While there are some exceptions to this (Weaver; EZH2 A677T), it highlights a potential link between these distinct rare syndromes that incomplete disruption of PRC2 binding and H3K27me3 levels may be a common feature. Identifying common features such as this may have a major impact in the rare disease space as interest groups aim to find commonalities and use these to drive shared therapeutic discovery efforts.

## Discussion

The mechanisms we have uncovered for how ASXL1 truncation impacts H2AK119ub1 and the Polycomb repressive system are integral to understanding the pathogenesis of several human diseases. This includes Bohring-Opitz syndrome, and other ASXL-related disorders; Shashi-Pena and Bainbridge-Roper syndromes (Doyle *et al*., 2022b). Indeed 15-30% of patients with AML and other myeloid disorders feature truncating nonsense and frameshift mutations in ASXL1. Therefore, the mechanistic insights we have discovered are likely to be critical in the myeloid context (Carbuccia *et al*., 2009; Döhner *et al*., 2022; Gelsi-Boyer *et al*., 2009). AML frequently features mutually exclusive genetic events that impact PRC1 and PRC2, including recurrent BCOR (PRC1) and EZH2 (PRC2) mutations (Schaefer et al., 2022; Simon et al., 2012). These, and other chromatin regulator-related drivers, such as TET2 and DNMT3A mutations, may suggest a shared mechanism of Polycomb repression dysfunction across these patients.

Our work highlights the importance of endogenous systems to understand the role of disease-driving genes, as key molecular effects may be missed due to technical artefacts of exogenous expression systems. Particularly in the case ASXL1-3 proteins, which are unstable and difficult to detect through antibody-dependent assays. The observation of the high instability of wild-type ASXL1 illustrates the critical nature of Polycomb protein dosage for maintaining a correct balance of repressive modifications. Furthermore, the discovery that imbalances of PR-DUB stoichiometry can impact its catalytic activity mirrors similar rheostat-like combinatorial assemblies of PRC2 and PRC1 complexes with different catalytic potencies displaying evidence of sub-functionalisation in certain cellular contexts, such as X-chromosome inactivation and Polycomb body formation (Almeida *et al*., 2017; Fursova *et al*., 2019; Kundu et al., 2017; Lavarone *et al*., 2019; Margueron *et al*., 2008; Scelfo et al., 2019). Interestingly, we found no evidence for changes in the stability of BAP1 or allosteric activation of ASXL1 through DEUBAD mono-ubiquitination, with evidence instead suggesting that a switch in the ASXL-binding partner of BAP1 may be sufficient to drive changes in catalytic activity.

Our analysis contrasting the impacts on Polycomb repression across genetically diverse chromatinopathy models (Borges *et al*., 2026; Conway *et al*., 2021; Deevy *et al*., 2025) highlights an emerging theme – that imbalances in H2AK119ub1 can play a major role in driving human disease, alongside the established roles of H3K27me3 loss of equilibrium in cancer (Conway et al., 2015; Tamburri *et al*., 2022). Our work and others’ highlights that the PR-DUB complex is at the forefront of retaining this balance in both H2AK119ub1 levels and spatial distribution in order to maintain plasticity of cell fate. This is further exemplified by the mutually exclusive occurrence of BAP1 heterozygous loss and ASXL1 truncating mutations in AML (Andricovich et al., 2025; Döhner *et al*., 2022), as either event alone can drive these H2AK119ub1 imbalances in divergent directions, with a potentially convergent impact on PRC2 activity. Indeed, several rare monogenic syndromes have been identified that are defined by variants in the H2AK119ub1 readers – JARID2, RYBP and DNMT3A (Deevy and Bracken, 2019; Verberne et al., 2021; Weinberg *et al*., 2021; Weisz-Hubshman et al., 2025) – alluding to the importance of the modification in these systems as well.

From a clinical standpoint, the discovery that reductions in PRC2 catalytic activity may be a common feature across several Polycomb-driven chromatinopathies – Weaver (EZH2), Bohring-Opitz (ASXL1), Luo-Schoch-Yamamoto (RING1A/B) and Kury-Isidor (BAP1) syndromes – may prove to be highly impactful to direct the discovery and implementation of therapeutics beyond gene therapies. Syndromes defined by PRC1, PRC2 and PR-DUB, such as Cohen-Gibson (EED) and Imagawa-Matsumoto (SUZ12) syndrome, may share these mechanistic changes to the epigenome and plasticity (Deevy and Bracken, 2019; Doyle *et al*., 2022b; Tamburri *et al*., 2022). Due to the pleiotropic nature of many chromatinopathies across diverse organ systems, individual gene therapies are difficult to engineer and successfully deliver to the necessary cell types. Finding common ground across these syndromes holds promise for the utilisation or development of epigenetic targeted compounds to restore equilibrium in these key repressive histone modifications. For instance, KDM6A/B inhibition has shown promising *in vitro* efficacy to increase H3K27me3 in Weavers syndrome models (Gao et al., 2024), while DOT1L inhibition may be a means to drive spatially controlled increases in PRC1 catalytic activity (Neville et al., 2026). These avenues may prove beneficial options for chromatinopathy therapeutics, along with other pharmacological approaches that can impact Polycomb related histone modifications such as BAP1, NSD2 and BAF complex targeting compounds (Hanley et al., 2023; Kofink et al., 2022; Wang *et al*., 2021) as these proteins and their complexes function to oppose Polycomb activities (Kadoch et al., 2017; Streubel *et al*., 2018).

### Limitations of study

While we have comprehensively characterised the mouse *ASXL1* G643Wfs variant – homologous to human ASXL1 G646Wfs – there are over one hundred syndromic ASXL frameshift and nonsense variants that we have not tested. The majority of ASXL1 variants occur in the final exon, and, therefore potentially escape nonsense-mediated decay like G643fs. However, some ASXL1, ASXL2 and many ASXL3 variants occur in the second to last exon (Bainbridge *et al*., 2013; McGrath *et al*., 2022). Therefore, future studies should characterise the broad scope of ASXL – syndrome variants for their impact – whether they drive ASXL stability or indeed haploinsufficiency. A comprehensive atlas of any distinct impacts could serve a critical role for determining clinical outcomes, biomarker interpretation, and future treatment selection.

## Resource Availability

### Lead contact

Requests for further information and resources should be directed to and will be fulfilled by the lead contact, Dr Eric Conway

### Materials availability

Reagents generated for this study will be made available upon reasonable request.

### Data and code availability

The sequencing data have been deposited into the Gene Expression Omnibus (GEO) repository.

- ChIP-seq data accession number: GSE337465
- RNA-seq data accession number: GSE337436

The mass spectrometry proteomics data have been deposited to the ProteomeXchange Consortium via the PRIDE partner repository with the dataset identifier PXD080492. No original code was generated for this study.

## Acknowledgements

We thank members of the Conway and Coughlan labs for helpful discussion and critical reading of the manuscript. We would like to thank the ASXL Rare Research Endowment foundation research and family community, for helpful discussions and feedback. We are grateful to the Genomics Core at University College Dublin Conway Institute and the IEO Genomics Unit for expertise and assistance with next-generation sequencing. We are grateful to the Trinity St. James Hospital Haematology Biobank for their support. We are especially thankful to the patients who agreed to provide primary BMNC to the biobank for their contribution. We thank Dr Daniel Fernandez-Perez for helpful discussions regarding computational analyses. Work in the Conway lab was supported by a Wellcome Trust Early Career Award [225152/Z/22/Z], a Worldwide Cancer Research award (23-008) and a Research Ireland Pathway Program grant (21/PATH-S/9384). E.J.D. was supported by a PhD fellowship from the Research Ireland Government of Ireland Postgraduate Scholarship Programme (GOIPG/2023/3637). S.B. was supported by a PhD fellowship from the Research Ireland Government of Ireland Postgraduate Scholarship Programme (GOPIG/2024/3239). M.B. was supported by a University College Dublin School of Biomolecular and Biomedical Sciences Research Scholarship. Work in the Pasini lab was supported by the Italian Association for Cancer Research, AIRC (IG-2022-27694), and by Worldwide Cancer Research (22-0027). Work in the Coughlan lab was supported by a Research Ireland Pathway Program grant (22/PATH-S/10642) and UCD Ad Astra Start-Up.

## Author contributions

E.C conceived the study. E.J.D and E.C designed the experiments. E.J.D performed the majority of the experiments (including genome editing, ChIP-seq, RNA-seq, co-IP) and analyses (RNA-seq and ChIP-seq). M.B, E.J.D and A.Y.C optimised CUT&Tag protocol for primary patient sample use. M.B, T.S, A.M.M and N.O planned and performed the primary AML sample isolation and subsequent CUT&Tag experiments and analysis. S.B supported the wet-lab experiments. D.P and M.B support genome mapping experiments. E.J.D and E.D performed and analysed the proteomics experiments. E.J.D and E.C wrote the manuscript and edited the figures.

## Declaration of interests

The authors declare no competing interests.

### Experimental Model and Subject Details Cell culture

Mouse embryonic stem cells (mESCs) were grown on 0.1% gelatin-coated culture dishes in Glasgow minimum essential medium (Sigma) supplemented with 20% (v/v) heat-inactivated FBS (Gibco), 100 U/mL penicillin/ 100 µg/mL streptomycin (Gibco), 50 µM β-mercaptoethanol, 1:100 GlutaMAX (Gibco), 1:100 non-essential amino acids (Gibco), 1 mM sodium pyruvate (Gibco), 1:500 leukemia inhibitory factor (LIF; produced in-house), 3 µM CHIR99021 (Cayman Chemical Company), and 1 µM PD0325901 (Cayman Chemical Company). HEK293T cells and NTERA-2 cells (for ChIP-Rx spike-in) were grown on TC-treated culture dishes in high-glucose Dulbecco’s modified Eagle’s medium (Sigma) supplemented with 10% (v/v) FBS (Gibco) and 100 U/mL penicillin/100 µg/mL streptomycin (Gibco).

### AML patient bone marrow mononuclear cell isolation

This study was approved by the St James’s Hospital and Tallaght Hospital Joint Research Ethics Committee. Following written informed consent, bone marrow aspirates were obtained from patients with AML presenting at St James’s Hospital, Dublin. Bone marrow mononuclear cells were isolated from aspirates by density-gradient centrifugation and cryopreserved according to modified published protocols (Brophy *et al*., 2023), and stored in the Trinity St James’s Haematology Biobank until required.

## Method Details

### CRISPR genome editing

sgRNA targeting the 3’ UTR of *ASXL1* were cloned into pSpCas9(BB)-2A-GFP (PX458) while to induce single strand DNA breaks at 3’ end, or in the final exon, of *ASXL1* gene, sgRNAs targeting the last exon of ASXL1 were cloned into the Cas9 nickase expression plasmid pSpCas9n(BB)-2A-GFP (PX461) (Addgene #48140). For homology directed repair (HDR) template, gBlocks for FLAG/HA-ASXL1 WT and FLAG/HA-ASXL1 G643Wfs repair templates were purchased (IDT) and cloned into the pCR8/GW/TOPO Gateway entry vector (Invitrogen). HDR templates contained ∼800bp homology arms, 2x FLAG, 3x HA, T2A and Blasticidin resistance gene along with single nucleotide edits to the PAM site to prevent allele re-targeting. E14 mouse embryonic stem cells were transfected with 3μg of sgRNA and 7μg of HDR template in pCR8 using Opti-MEM and the Lipofectamine 2000 reagent, as per manufacturer’s instructions (ThermoFisher). Media was changed 24 hours after transfection. 48 hours after transfection, cells were prepared for sorting in PBS, 1% FBS and 2mM EDTA and filtered at 70μm. Cells were sorted to isolate GFP high populations using a CytoFLEX SRT flow cytometer. Cells were seeded at low density and grown for 10-14 days. Individual clones were picked and dissociated in 2.5% trypsin, neutralised and plated. Clones were screened at the gDNA level for the desired heterozygous knock-in allele, and un-edited wild-type allele, via PCR and Sanger Sequencing (Eurofins). BAP1 knockout cells were generated previously (Conway *et al*., 2021).

### Plasmids

The full-length ASXL1 ORF was cloned via PCR amplification of 3 segments, before Gibson assembly into pCR8/GW/TOPO plasmid. ASXL1 404*, 635*, 646*, 693*, 733* and 1028* fragments were PCR amplified from pCR8 ASXL1 WT. PCR products were inserted into the pCR8/GW/TOPO Gateway cloning entry vector (Invitrogen). Coding sequences were subcloned into Gateway compatible expression vectors (pLEX_305-3xHA or pCAG-2xFLAG-HA) using LR clonase II (Invitrogen). ASXL1 K351R mutants were introduced using Q5 site-directed mutagenesis (NEB) according to manufacturer’s instructions. pCAG-2xFLAG/HA-BAP1 was generated previously (Conway *et al*., 2021).

### Western blot

Whole-cell protein samples were prepared in ice-cold high-salt buffer (50 mM Tris-HCl at pH 7.2, 300 mM NaCl, 0.5% [v/v] NP-40, 1 mM EDTA at pH 7.4, 2 µg/ mL aprotinin, 1 µg/mL leupeptin, 1 mM PMSF). Cell suspensions were sonicated and then rotated for 20 min at 4°C to ensure sufficient lysis. Lysates were next clarified by centrifugation at ≥20,000 RCF for 20 min at 4°C. Normalized protein lysates were denatured and separated on Bolt Bis-Tris (Invitrogen) gels and then transferred to 0.2 µM nitrocellulose membranes (Amersham). Membranes were blocked against nonspecific binding and then incubated with the relevant primary antibodies overnight at 4°C and secondary antibodies for 1 hour at room temperature. Relative protein levels were determined by chemiluminescence or infrared fluorescence detection on a LI-COR Odyssey Fc or CLx. Primary antibodies used for western blot were: FLAG (F1804; Merck), HA (C29F4; Cell Signaling Technology), BAP1 (D7W7O; Cell Signaling Technology), H2AK119ub1 (D27C4; Cell Signaling Technology), H3 (ab1791; Abcam). Secondary Antibodies used for western blot were: Anti-mouse/ anti-rabbit HRP (7076/ 7074S; Cell Signaling Technology), Anti-mouse 680CW/Anti Rabbit 700CW (926-68070/926-32211; Licor).

### Co-immunoprecipitation

Mouse embryonic stem cells were harvested by centrifugation, washed twice with ice cold PBS, and resuspended in Buffer A (25mM HEPES pH 7.6, 5mM MgCl_2_, 25mM KCl, 0.05mM EDTA, 10% (v/v) glycerol, 0.1% NP-40, 1mM DTT, combined with protease inhibitors) and incubated with rotation at 4°C for 10 minutes. Cell nuclei were pelleted by centrifugation at 500rcf at 4°C for 10 minutes. Nuclei were lysed in Buffer C (10mM HEPES pH 7.6, 3mM MgCl_2_, 100mM KCl, 0.5mM EDTA, 10% (v/v) glycerol, 1mM DTT, combined with protease inhibitors). Then, 300mM (NH_4_)_2_SO4 dissolved in Buffer C was added to the samples which were incubated on ice for 20 minutes. Samples were ultra-centrifuged at 350,000rcf for 15 minutes at 4°C using an SW 55 Ti swinging-bucket rotor in a Beckman Coulter Optima L-100XP. The supernatant was collected, and nuclear extracts were precipitated by adding 300mg (NH_4_)_2_SO4 per 1ml of sample. The samples were incubated on ice for 20 minutes before repeating ultracentrifugation using the same parameters as above. Nuclear pellets were resuspended in IP buffer (300mM NaCl, 50mM Tris-HCl pH 7.5, 1mM EDTA, 1% (v/v) Triton-X100, 1mM DTT, combined with protease inhibitors) and protein quantification and sample normalization was carried out using the Bradford assay. For each co-IP, 1μl Benzonase Nuclease (Sigma-Aldrich) and 1μg of the respective antibody was added to 1mg of nuclear protein. Samples were rotated at 4°C overnight. The next day, washed Protein G Dynabeads (Invitrogen) were added to each sample followed by rotation at 4°C for 2 hours. Beads were then washed four times with IP buffer followed by two cold PBS washes. Beads were stored at -80°C until further processing.

### Mass spectrometry

Beads from the co-immunoprecipitation were washed with phosphate buffered saline and 8M urea was added to a volume of 200μl. Following this, 5mM DTT was added to the beads and incubated at 60°C for 30 minutes. Then, 9mM iodoacetamide solution was added to the beads and incubated in the dark at room temperature for 30 minutes. 2.2mM CaCl_2_ was added to the beads followed by addition of 12.8mM ammonium bicarbonate pH 8. Following this, 1μg of sequencing grade trypsin (Sigma-Aldrich) was added to each reaction and incubated at 37°C overnight on a thermomixer at 350rpm. The next morning, 1% trifluoroacetic acid was added to stop the reaction. The tryptic digests were then desalted using Pierce C18 Spin Tips (Thermo Scientific) according to the manufacturer’s protocol. Peptides (500-1000ng) were loaded onto Evotips as per manufacturer’s instructions (Evosep). Briefly, Evotips were activated by soaking them in isopropanol, primed with 20 μL buffer B (ACN, 0.1% FA) by centrifugation for 1 min at 700 g. Tips were soaked in isopropanol and equilibrated with 20 μL buffer A (MS grade water, 0.1% FA) by centrifugation. Another 20 μL buffer A was loaded onto the tips and the samples were added on top of that. Tips were centrifuged and washed with 20 μl buffer A followed by overlaying the C18 material in the tips with 100 μL buffer A and a short 20 s spin. The samples were analysed by the Mass Spectrometry Resource (MSR) in University College Dublin on a Bruker TimsTOF Pro mass spectrometer connected to a Evosep One chromatography system. Peptides were separated on an 8 cm analytical C18 column (Evosep, 3 μm beads, 100 μm ID) using the pre-set 30 samples per day gradient on the Evosep one. The Bruker TimsTOF Pro mass spectrometer was operated in positive ion polarity with TIMS (Trapped Ion Mobility Spectrometry) and PASEF (Parallel Accumulation Serial Fragmentation) modes enabled. The accumulation and ramp times for the TIMS were both set to 100 ms., with an ion mobility (1/k0) range from 0.62 to 1.46 Vs/cm. Spectra were recorded in the mass range from 100 to 1,700 m/z. The precursor (MS) Intensity Threshold was set to 2,500 and the precursor Target Intensity set to 20,000. Each PASEF cycle consisted of one MS ramp for precursor detection followed by 10 PASEF MS/MS ramps, with a total cycle time of 1.16 s.

### ChIP-Rx

ChIP experiments were performed according to protocols as described previously (Ferrari et al., 2014). For all ChIPs, 1% formaldehyde cross-linked chromatin was sheared to 500–1000 bp fragments by sonication. Chromatin was then incubated overnight in IP buffer (33 mM Tris-HCl pH 8, 100 mM NaCl, 5 mM EDTA, 0.2% NaN3, 0.33% SDS, 1.66% Triton X-100) at 4°C with the indicated antibodies. 5μg antibodies were used per 500μg chromatin for RING1B (D22F2; Cell Signaling Technology) and SUZ12 (D39F6, Cell Signaling Technology) ChIPs. 3μg antibodies per 250μg of chromatin were used for histone modification ChIPs, H2AK119ub1 (D27C4; Cell Signaling Technology), H3K27me3 (C36B11, Cell Signaling Technology). All ChIPs were supplemented with 5% spike-in of NTERA-2 human chromatin (prepared in the same manner). The next day, chromatin lysates were incubated for 3 hours with protein-G Sepharose beads (GE Healthcare). Beads were washed 3 × with low-salt buffer (150mM NaCl, 20mM Tris-HCl pH 8, 2mM EDTA, 0.1% SDS, 1% Triton X-100) and 1 × with high-salt buffer (500mM NaCl, 20mM Tris-HCl pH 8, 2mM EDTA, 0.1% SDS, 1% Triton X-100), and then re-suspended in de-crosslinking solution (0.1M NaHCO3, 1% SDS). DNA was purified with QIAquick PCR purification kit (QIAGEN) according to manufacturer’s instructions. DNA libraries were prepared with 2–10 ng of DNA using an in-house protocol (Blecher-Gonen et al., 2013) by the IEO genomic facility and sequenced on an Illumina NovoSeq 6000. ChIP-Rx experiments were performed in biological duplicate and includes two distinct ASXL1 BOS clones.

### RNA-seq

Total RNA from three biological replicates was isolated from ESCs using the NucleoSpin mini kit for RNA purification (Macherey-Nagel), according to manufacturer’s instructions. RNAseq libraries were prepared by Novogene as follows; mRNA was purified from total RNA using poly-T oligo-attached magnetic beads. Following fragmentation, first strand of cDNA was synthesized using random hexamers, followed by second strand 72 synthesis using dTTP. cDNA was subject to end repair, A-tailing, adapter ligation before being size selected and amplified for final library generation. Library was assessed by Qubit, Bioanalyser and qRT-PCR for quality control and quantification. Libraries were sequenced paired end to an approximate read depth of 55 million paired end reads on Illumina NovaSeqX plus platform.

### CUT&Tag

CUT&Tag was performed on cryopreserved patient-derived BMNCs following modified published protocols (Kaya-Okur et al., 2019).

For each reaction, 90,000 freshly thawed patient cells were used together with 10,000 mouse embryonic stem cells as spike-in controls. Each experiment was performed with technical duplicates. Cells were crosslinked in 0.1% formaldehyde in PBS for 1 minute before being quenched with 125mM glycine for 5 minutes. Nuclei were obtained by incubation with nuclei extraction buffer (20mM HEPES–KOH pH 7.9, 10 mM KCl, 0.1% Triton X-100, 20% glycerol, 0.5 mM spermidine, 1x protease inhibitor cocktail) for 10 minutes on ice. Nuclei were bound to activated concanavalin-A beads (Cell Signaling) and subsequently incubated with 1ug of primary antibody, H2AK119ub1 (D27C4; Cell Signaling Technology) and H3K27me3 (C36B11, Cell Signaling Technology), in Antibody150 buffer (20 mM HEPES pH 7.9, 150 mM NaCl, 0.01% digitonin, 2 mM EDTA, 0.5 mM spermidine, 1x protease inhibitor cocktail) overnight at 4°C with rotation.

The following day, samples were incubated with anti-rabbit IgG secondary antibody (EpiCypher) for 30 minutes at room temperature. After washing in Digitonin150 buffer (20 mM HEPES pH 7.9, 150 mM NaCl, 0.01% digitonin, 0.5 mM spermidine, protease inhibitors), samples were resuspended in pAG-Tn5 (Epicypher) in Digitonin300 buffer (20 mM HEPES pH 7.9, 300 mM NaCl, 0.01% digitonin, 0.5 mM spermidine, protease inhibitors) and rotated for 1 hour at room temperature. Tagmentation was performed by incubating samples in tagmentation buffer (20 mM HEPES pH 7.9, 300 mM NaCl, 0.01% digitonin, 10 mM MgCl_2_, 0.5 mM spermidine, protease inhibitors) at 37°C for 1 h. Beads were then washed in TAPS buffer (10 mM TAPS, 0.2 mM EDTA), and incubated with SDS release buffer (10 mM TAPS, 0.1% SDS) for 1 hour at 58°C, followed by SDS quenching (0.67% Triton X-100).

Libraries were PCR amplified directly from bead-bound material using custom indexing primers and polymerase (NEB). PCR conditions were: 5 min at 58°C, 5 min at 72°C, 45 s at 98°C, followed by 21 cycles of amplification 98°C for 15 s, 60°C for 10 s, with a final extension at 72°C for 1 minute. PCR products were purified using 1.3X AMPure beads (Beckman Coulter) and eluted in 0.1X TE.

Libraries were quantified using a Qubit fluorometer, and library distribution was assessed on a TapeStation prior to sequencing. Libraries were sequenced as 50 bp paired-end reads on the NextSeq 2000, with raw sequencing depths of ∼10m read pairs.

### Data analysis

#### ChIP-seq analysis

Paired-end DNA reads were processed through fastp to trim adapters and to remove low quality nucleotides at read ends (Chen et al., 2018). Quality-filtered DNA reads were aligned to the mouse reference genome, mm10, and human reference genome (hg38) using Bowtie v1.2.2 with the parameters -I 10, - X 1000 (Langmead et al., 2009). Reads mapped to both mm10 and hg38 were discarded.

The intensity of histone modifications or transcription factor binding was represented through boxplots or heatmaps, both generated from BigWig files that were obtained using the function bamCompare from deepTools 3.1 (Ramirez et al., 2016) with parameters –binSize 50 –extendReads. The – scaleFactors parameter of bamCompare was set to (1/total mapped reads)∗1,000,000 to normalize for the differences in sample library size. Samples obtained with the ChIP-Rx method, were normalized using (1/hg38 mapped reads)∗1,000,000 as –scaleFactors parameter (Orlando et al., 2014). The preparation of the heatmaps required the generation of a data matrix through computeMatrix with parameters – referencePoint TSS/center -a 5000 -b 5000 (Ramirez *et al*., 2016). The data matrix was converted into a heatmap by plotHeatmap (Ramirez *et al*., 2016). Boxplots were prepared using multiBigwigSummary in BED-file mode and using as bed file the regions corresponding to promoters (TSS ± 2.5 kbp) or intergenic regions (Ramirez *et al*., 2016). Intergenic sites were defined by subtracting from the mouse genome, the region included between the transcription start and end sites of each gene. P values and effect sizes displayed in boxplot quantifications of ChIP-seq data were calculated using non-parametric rank tests. For p values, Mann-Whitney’s U Test was performed, meanwhile for the effect sizes we applied the rank-biserial correlation. These calculations were done using the r functions wilcox.test (r-base) and rank_biserial (effectsize package) (Sullivan and Feinn, 2012), respectively.

Chromosome-wide plots were generated using MultiBigWigSummary in BED-file mode, with the BED file containing chromosomes separated into 25kb bins and plots were generated using the ggplot2 package in R .

#### RNA-seq analysis

Low quality reads were filtered and then remaining reads were aligned to the mm10 reference genome using HISAT2 (Mortazavi et al., 2008). The featureCounts tool from the Subread package was used to calculate read counts from which FPKM was then calculated, accounting for sequencing depth and gene length (Liao et al., 2013). Differentially expressed transcripts were identified using the DESeq2 R package (Love et al., 2014), applying a fold change threshold of >1 for upregulated genes and a Benjamini-Hochberg-adjusted *P*-value of < 0.05. Gene Ontology (GO) Enrichment analysis was performed using the clusterProfiler R package (Yu et al., 2012), with significance thresholds of *P*-value < 0.05 and *q*-value < 0.05. Three replicate FPKM values were averaged for each cell line, log2-transformed and Zscore normalisation was applied. Heatmaps were generated using the Heatmap function (base R). Unsupervised hierarchical clustering was applied using Pearson correlation distance and ward.D2. Venn Diagrams were generated using the eulerr package (Larsson and Gustafsson, 2018).

#### Proteomics analysis

Bruker mass spectrometric data from the Tims-Tof was processed using the MaxQuant (version 2.0.3.0) incorporating the Andromeda search engine. To identify peptides and proteins, MS/MS spectra were matched against a mus musculus uniport database containing 82,427 entries (release 2023_03). All searches were performed using the default setting of MaxQuant, with trypsin as the specified enzyme allowing two missed cleavages and a false discovery rate of 1% on the peptide and protein level. The database searches were performed with carbidomethyl as a fixed modification and acetylation (protein N terminus) and oxidation (M) as variable modifications. For the generation of label free quantitative (LFQ) ion intensities for protein profiles, signals of corresponding peptides in different nano-HPLC MS/MS runs were matched by MaxQuant in a maximum time window of 1 min. The ‘proteinGroups.txt’ output file was filtered for contaminants and reverse hits using Perseus version 2.0.10.0. Volcano plots were generated using Perseus version 2.0.10.0. For stoichiometric analysis, iBAQ values for IgG control were first subtracted from each experimental replicate. The subsequent iBAQ value of each identified protein was normalised to the respective bait protein in each IP. The stoichiometry values were averaged across replicates, reported as the mean. Plots were formatted using GraphPad PRISM version 10.5.0.

#### CUT&Tag analysis

Paired-end DNA reads were processed using fastp (v0.23.4) (Chen *et al*., 2018) (Chen et al., 2018) to remove low-quality reads. Filtered reads were aligned to the hg38 and mm10 reference genomes using Bowtie2 (v2.3.4.1) (Langmead *et al*., 2009) with parameters -I 10 -X 700. Duplicate reads mapping to both human and mouse genomes were discarded before PCR duplicates were removed with sambamba (v1.0.1). Resulting BAM files were used to generate normalised coverage tracks with deeptools (v3.5.6) bamCoverage (Ramirez *et al*., 2016) using (1e6 / Number of spike in mapped reads) to normalise. Heatmaps were prepared using deeptools computeMatrix with parameters –referencePoint TSS -a 5000 -b 5000 --missingDataAsZero and plotted with plotHeatmap.

---

**Figure S1.**
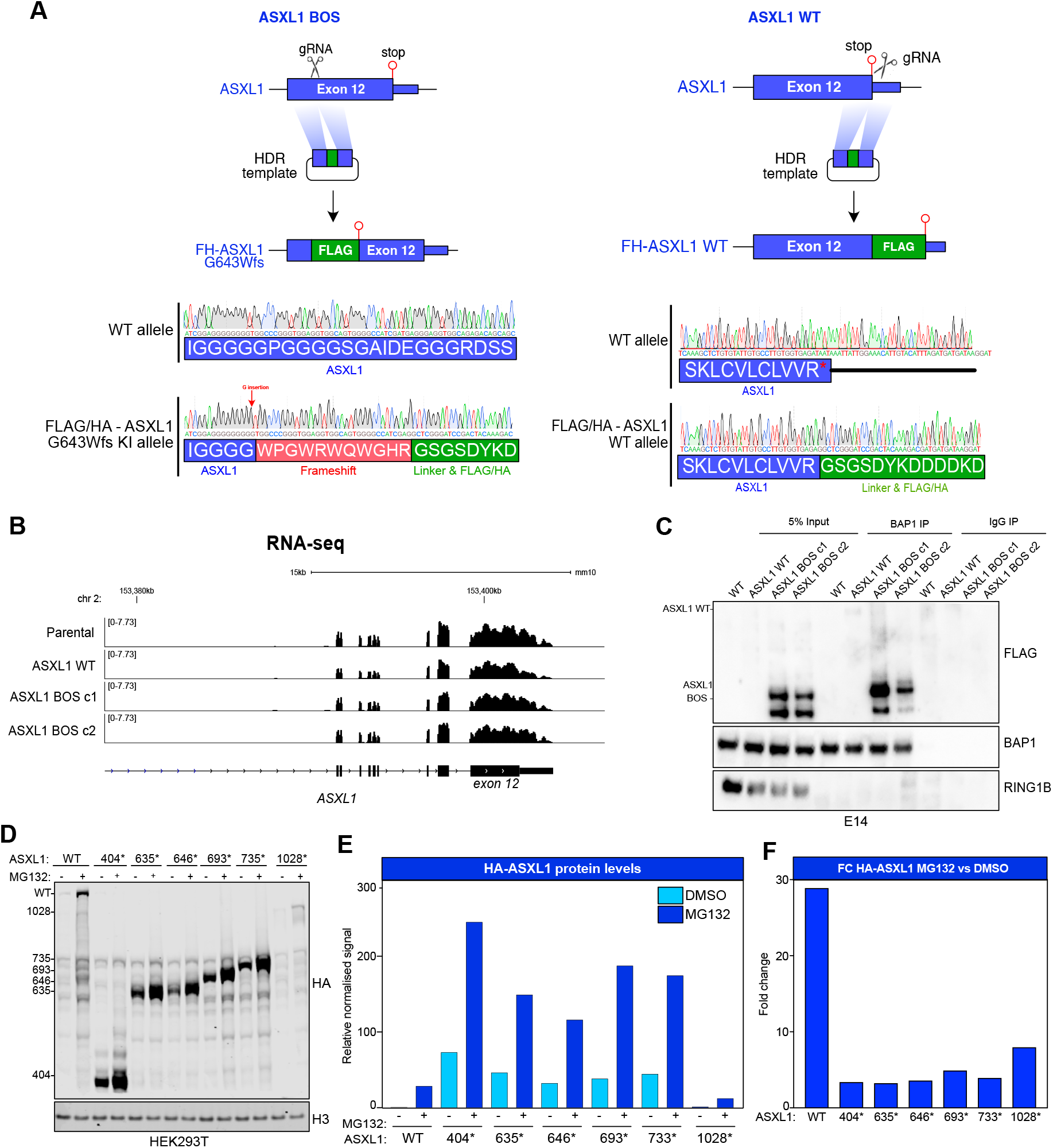
Related to Figure 1. A. Top. Detailed schematic of CRISPR nickase knock-in strategy to develop tagged heterozygous *ASXL1*^*FLAG-G643fs/+*^ (ASXL1 BOS) and *ASXL1*^*FLAG-WT/+*^ (ASXL1 WT) embryonic stem cells line. Bottom. Sanger sequencing traces of KI allele from ASXL1 BOS and ASXL1 WT mESC lines. B. Genome browser snapshot of RNA-seq at *ASXL1* gene locus in the indicated cell lines. C. Western blot with indicated antibodies on nuclear lysates from the indicated mESCs line following IgG or BAP1 endogenous co-immunoprecipitation. D. Western blot with indicated antibodies on whole cell lysates from HEK293T cells transiently transfected with the indicated plasmid construct followed by vehicle or MG132 treatment for 6 hours. E. Quantification of fluorescence-based western blots in panel D. HA signal is normalised relative to H3 loading control. F. Fold change in signal of fluorescence-based western blots in panel D in MG132 vs DMSO treated cells.

**Figure S2.**
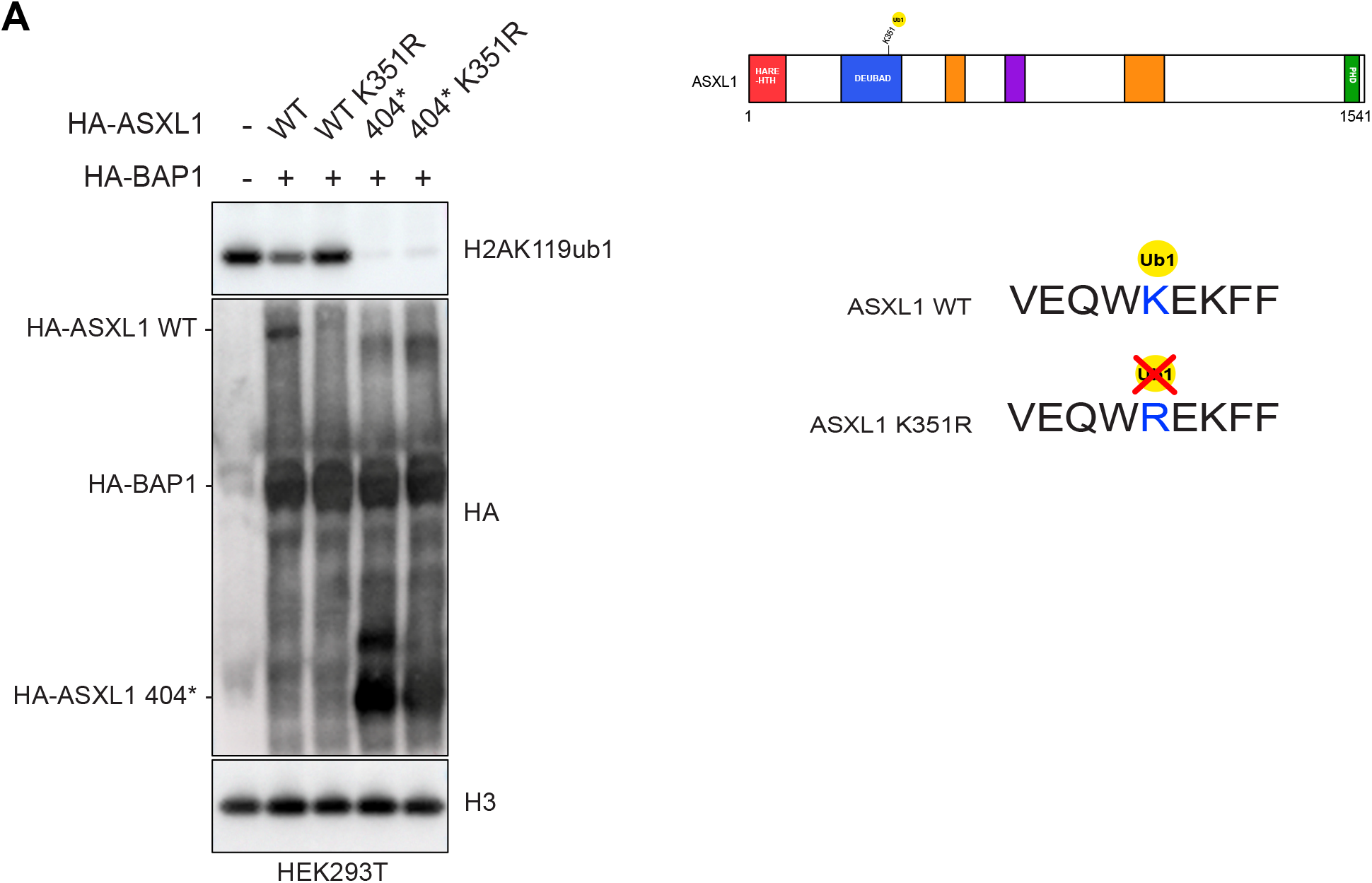
Related to Figure 2. A. Western blot with indicated antibodies on whole cell lysates of HEK293T cells following transient transfection with indicated plasmid construct.

**Figure S3.**
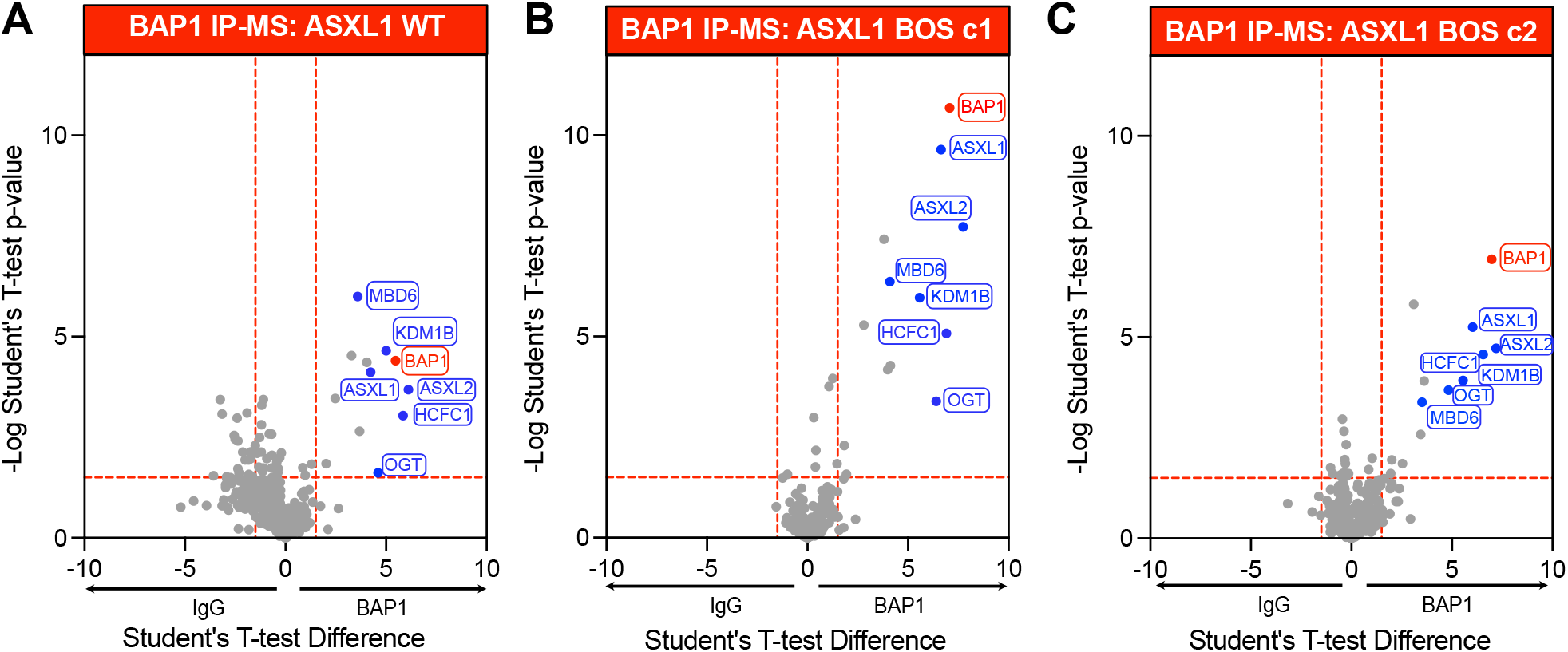
Related to Figure 3. A-C. Volcano plots of endogenous BAP1 IP mass spectrometry data relative to negative control IgG IP in ASXL1 WT (A), ASXL1 BOS c1 (B) and ASXL1 BOS c2 (C). T-test difference of LFQ values on the x-axis is plotted against -Log (T-test p-value). BAP1 (bait) is labelled in red and PR-DUB complex subunits are labelled in blue.

**Figure S4.**
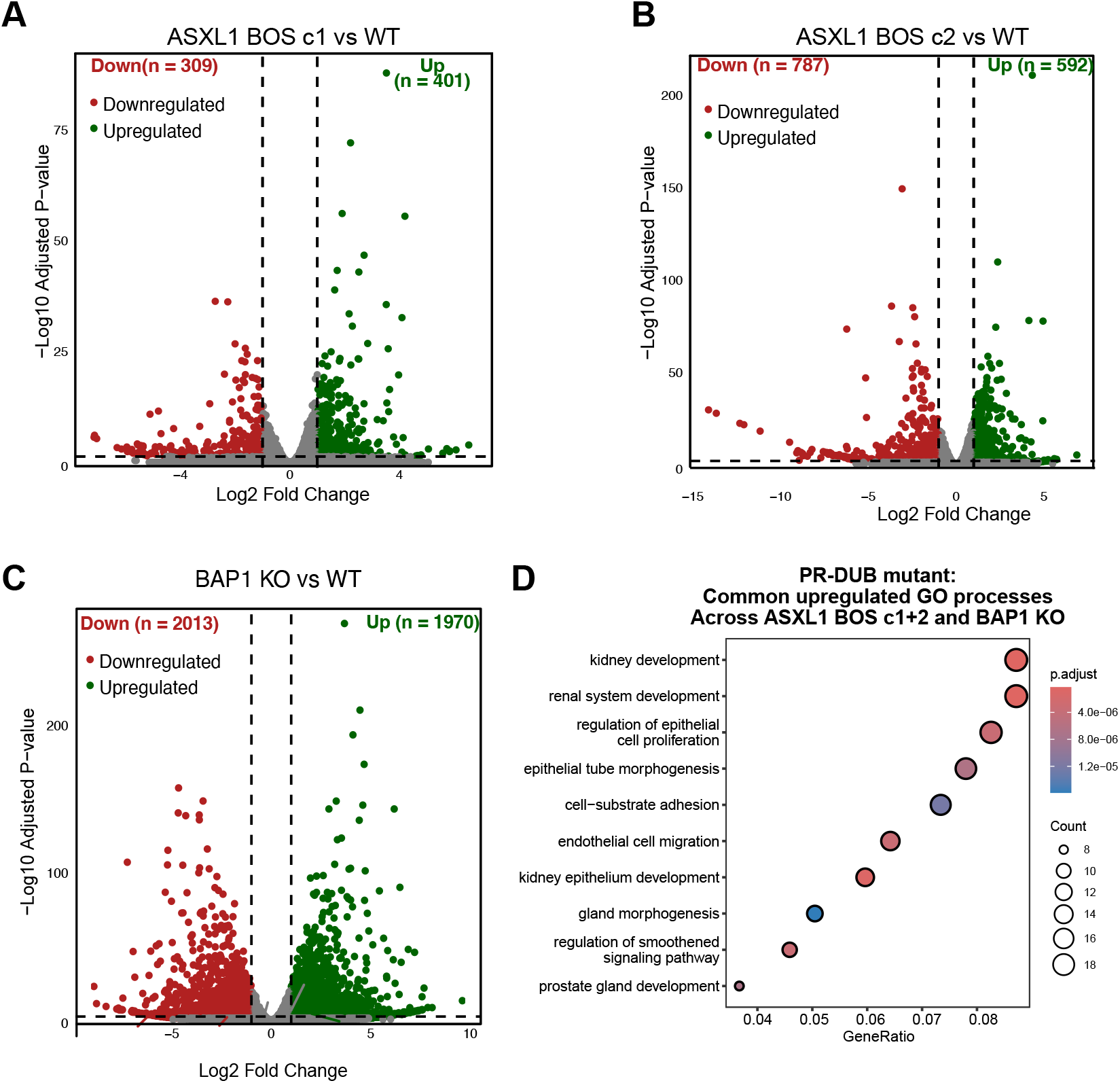
Related to Figure 5. A-C. Volcano plots of -log10 (p-value) against log2 fold change representing the differences in gene expression in the indicated cell lines vs WT ESCs. D. Gene ontology analysis of biological processes enriched in genes upregulated in both ASXL1 BOS clones and BAP1 KO mESCs.

